# Synchronous chromosome segregation in mouse oocytes is ensured by biphasic securin destruction and cyclin B1-Cdk1

**DOI:** 10.1101/824763

**Authors:** Christopher Thomas, Mark D. Levasseur, Rebecca J. Harris, Owen R. Davies, Suzanne Madgwick

## Abstract

Successful cell division relies on the faithful segregation of chromosomes. If chromosomes segregate prematurely the cell is at risk of aneuploidy. Alternatively, if cell division is attempted in the absence of complete chromosome segregation, non-segregated chromosomes can become trapped within the cleavage furrow and the cell can lose viability. Securin plays a key role in this process, acting as a pseudosubstrate to inhibit the protease separase that functions to cleave the cohesin rings that hold chromosomes together. Consequently, securin must be depleted ahead of anaphase, ensuring chromosome segregation occurs in time with the anaphase trigger. Here we find that MI mouse oocytes contain a large excess of securin over separase and reveal the existence of a novel mechanism that functions to promote the destruction of excess securin in prometaphase. Critically, this mechanism relies on key residues that are only exposed when securin is not bound to separase. We suggest that the majority of non-separase bound securin is removed by this mechanism, allowing for separase activity to be protected until just before anaphase. In addition, we further demonstrate the importance of complementary mechanisms of separase inhibition by directly measuring cleavage activity in live oocytes, confirming that both securin and inhibition by cyclin B1-Cdk1 are independently sufficient to prevent premature separase activation.

## Introduction

Successful cell division depends on the precise segregation of chromosomes into two equal sets. In mitosis and meiosis II this involves the segregation of sister chromatids. In meiosis I, pairs of homologous chromosomes are segregated in a reductional division that introduces genetic diversity. If chromosomes missegregate during these divisions, cells are at risk of aneuploidy, a hallmark of cancer and the primary genetic cause of miscarriage and developmental defects in babies^1–5^.

Through all cell divisions, the protease separase plays an essential role, cleaving the cohesin complexes that hold both sister chromatids and homologous chromosome pairs togethe^6,7^. It is therefore critical that separase remains inactive until all chromosomes are correctly aligned and the cell is prepared for anaphase.

Not surprisingly, separase activity is tightly regulated through the cell cycle. This regulation is executed through two distinct inhibitory pathways. In the primary pathway, separase directly interacts with securin, which acts as a pseudosubstrate inhibitor for separase^8^. In addition to this, upon phosphorylation by Cdk1, separase is inhibited by binding Cdk1’s activating partner cyclin B1^9–12^. The binding of separase with either securin or cyclin B1 is mutually exclusive and the relative contribution of each pathway varies depending on cell type and developmental state^10,12^. In female mouse meiosis II and eukaryotic mitosis, securin is largely responsible for separase inhibition, while primordial germ cells and early stage embryos rely primarily on cyclin B1-Cdk1-mediated separase inhibition^13–15^. Interestingly however, while securin binding to separase is the primary inhibitory mechanism in mitotic cells, it is also dispensable, demonstrating the compensatory nature of these two pathways^16–18^. Similarly, in fixed MI mouse oocytes Chiang et al. were able to inhibit securin or cyclin B1-Cdk1 mediated separase inhibition individually without observing segregation defects. Chromosome segregation errors were only detected when both inhibitory pathways were removed^19^.

Importantly, securin and cyclin B1 are not only involved in separase inhibition but their binding also plays a key role in priming separase for activation^20–22^. Furthermore, the degradation of securin and cyclin B1 must be coupled in order to ensure that separase activation and the poleward movement of chromosomes are timed correctly. In situations where these events become disconnected, anaphase is defective^23–25^.

In mitosis, the synchronous loss of cyclin B1 and securin is ensured by the similarity of their destruction mechanisms. Both are ubiquitylated by the Anaphase Promoting Complex/Cyclosome (APC/C) in metaphase^26,27^. Critically, this ubiquitylation relies on both the availability of the APC/C activator protein Cdc20 and on a short linear motif known as the D-box present in the N-terminus of both securin and cyclin B1^28–30^. Once all chromosomes are properly attached to spindle microtubules in metaphase, Cdc20 and the APC/C form a bipartite receptor for D-box docking, triggering securin and cyclin B1 destruction by the proteasome^31,32^. Prior to this, Cdc20 is sequestered by the spindle assembly checkpoint (SAC), a diffusible signal generated at each unattached kinetochore. The SAC functions to prevent docking of D-box motifs to the APC/C and thereby inhibit anaphase until all chromosomes are attached to spindle microtubules^33,34^.

In contrast, securin and cyclin B1 destruction in mouse oocyte meiosis I is initiated in prometaphase, prior to full chromosome alignment^35,36^. Initially this might seem like a failure to ensure accurate chromosome segregation. However, our recent work demonstrates that degradation of cyclin B1 in prometaphase is in fact a key feature of a mechanism that functions to prevent aneuploidy in mouse oocytes^36^. At this time point, destruction represents only the loss of non-Cdk1-bound cyclin B1. Here, cyclin B1 is targeted by an additional motif to the D-box, the PM motif. Importantly, the PM motif is only accessible in cyclin B1 that is not bound to Cdk1. By this strategy, an excess of cyclin B1 acts as an APC/C decoy to maintain Cdk1 activity and prolong prometaphase in oocyte meiosis I. This is critical in mouse oocytes since, in the absence of excess cyclin B1, spindle checkpoint activity is insufficient in preventing anaphase for long enough to fully align chromosomes, a process that takes hours rather than the tens of minutes to complete mitosis^36^.

Our discovery of a novel pathway of meiotic cyclin B1 regulation raises significant questions with regard to securin regulation. Since the D-box is insufficient for the correct degradation timing of cyclin B1 in oocytes, it seemed highly likely that additional pathways exist to degrade securin.

Here we find that securin exists in large excess to separase in oocyte MI. We identify key residues that function to promote the degradation of non-separase bound securin in prometaphase. Beyond this we show the importance of complementary mechanisms of separase inhibition in live oocytes.

## Results

### Securin destruction begins 2.5 hours ahead of separase activation in meiosis I in mouse oocytes

In mitosis, securin is destroyed alongside cyclin B1 and only in metaphase once the spindle checkpoint signals that all kinetochores are correctly attached to microtubules^26,27^. In contrast, while securin and cyclin B1 are also targeted simultaneously in meiosis I of mouse oocytes, their destruction is initiated much earlier, ahead of chromosome alignment and spindle migration in prometaphase (Fig. 1a). However, as division errors are rare in mouse oocytes, it seemed unlikely that this early targeting of securin affects separase activity so far ahead of anaphase.

**Figure 1.**
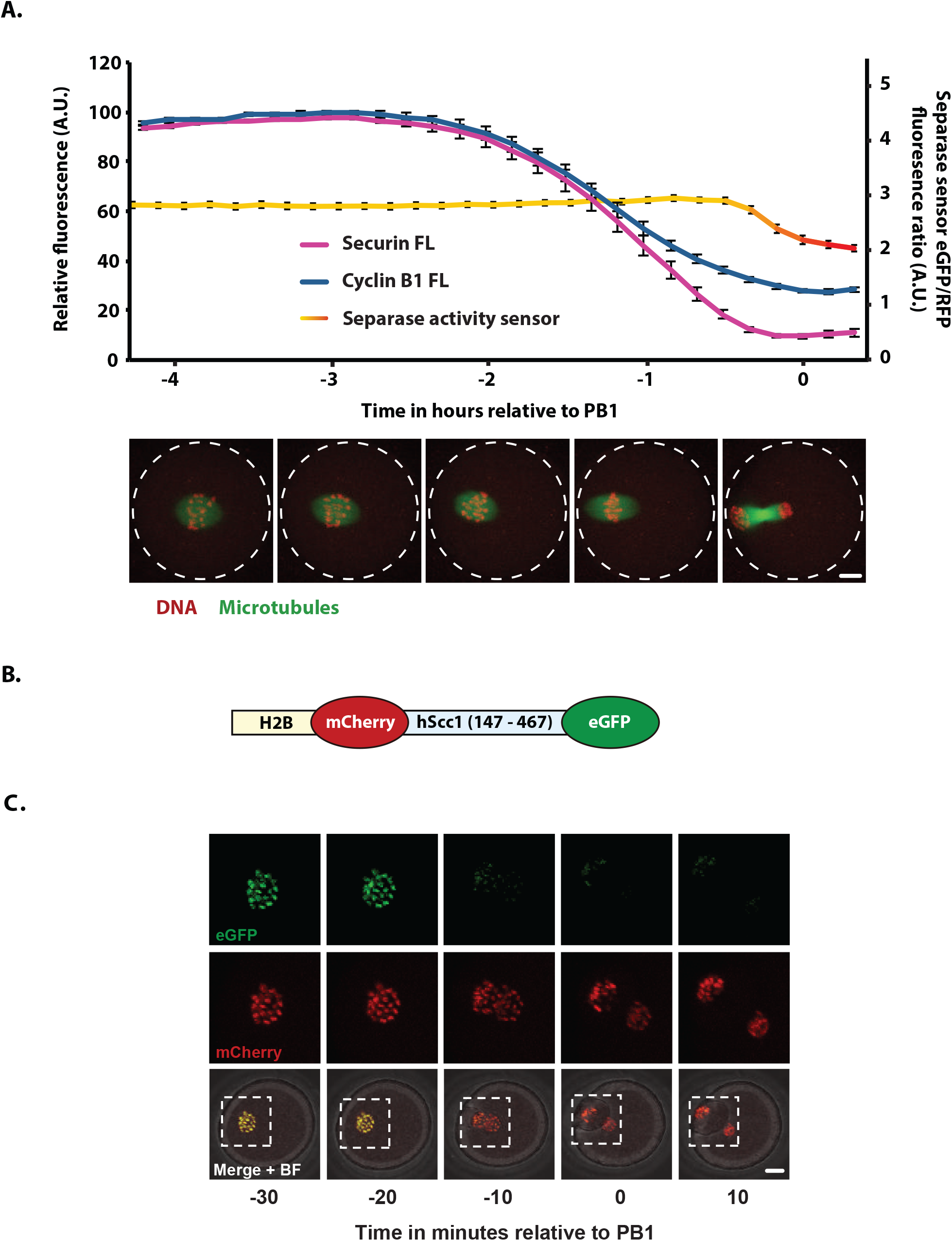
Securin destruction begins 2.5 hours ahead of separase activation in mouse oocyte meiosis I. (A) Graph showing the mean destruction profiles of VFP-tagged securin FL (magenta trace, n = 25) and cyclin B1 FL (blue trace, n = 62) alongside separase activity as determined by a separase activity biosensor (eGFP/RFP ratio, n = 20) in MI mouse oocytes relative to PB1 extrusion. Error bars ± SEM. Representative confocal images show a maturing oocyte expressing Map7-GFP (microtubules in green) and incubated with SiR-DNA (DNA in red) at times relative to the x-axis of the graph. Scale bar = 10 µm. (B) Schematic diagram of an H2B-mCherry-Scc1-eGFP separase activity biosensor. (C) Representative time-lapse images of an oocyte expressing the separase activity biosensor in Fig. 1B, imaged every 10 minutes at the times indicated relative to PB1 extrusion. eGFP (green), mCherry (red) and merged fluorescence + bright field (BF) images are shown. Scale bar = 10µm. eGFP and mCherry images correspond to the areas marked by the white dotted lines in the merge + BF images.

To assess whether degradation of securin and cyclin B1 in prometaphase caused separase activation, we used a separase activity biosensor generated by Nam et al. that has previously been validated in oocytes^37–39^. The sensor consists of nucleosome-targeted H2B protein fused to eGFP and mCherry fluorophores. Between the two fluorophores, an Scc1 peptide sequence is cleaved by active separase. Scc1 cleavage results in a yellow to red colour shift as the eGFP signal dissociates into the cytoplasm and mCherry remains bound to histones associated with chromosomal DNA. We injected germinal vesicle (GV) stage oocytes with mRNA encoding the biosensor and imaged the live cells through MI. A clear shift in colour was consistently observed just 20-30 minutes ahead of polar body extrusion (Fig. 1c). This timing was confirmed by the quantification of the eGFP/mCherry fluorescence ratio (Fig. 1a). The data indicate that separase is only active from 30 minutes ahead of polar body extrusion, with the majority of substrate cleavage taking place in the final 20 minutes. Importantly, separase becomes active more than 2 hours after the initiation of securin destruction.

### The D-box is insufficient for wild-type securin destruction in MI

In mouse oocytes, cyclin B1 is destroyed by a two-step mechanism; an initial period of prometaphase destruction requires both a D-box and an additional motif (the PM motif). This is followed by a second period of destruction as the D-box becomes sufficient once the SAC is satisfied in metaphase^36^. Given that securin and cyclin B1 destruction is synchronous in oocytes, we reasoned that securin may also be destroyed by a similar mechanism^35,36^. To test this, we initially designed and tested two fluorescent reporters; full-length securin (securin FL) and the N-terminal 101 residues (securin N101). Both securin reporters and all that follow were coupled to VFP to give a direct readout of exogenous protein level in the oocyte. Critically both constructs contain the D-box and all neighbouring lysine residues necessary for APC/C recognition and subsequent proteolysis (Fig. 2a).

**Figure 2.**
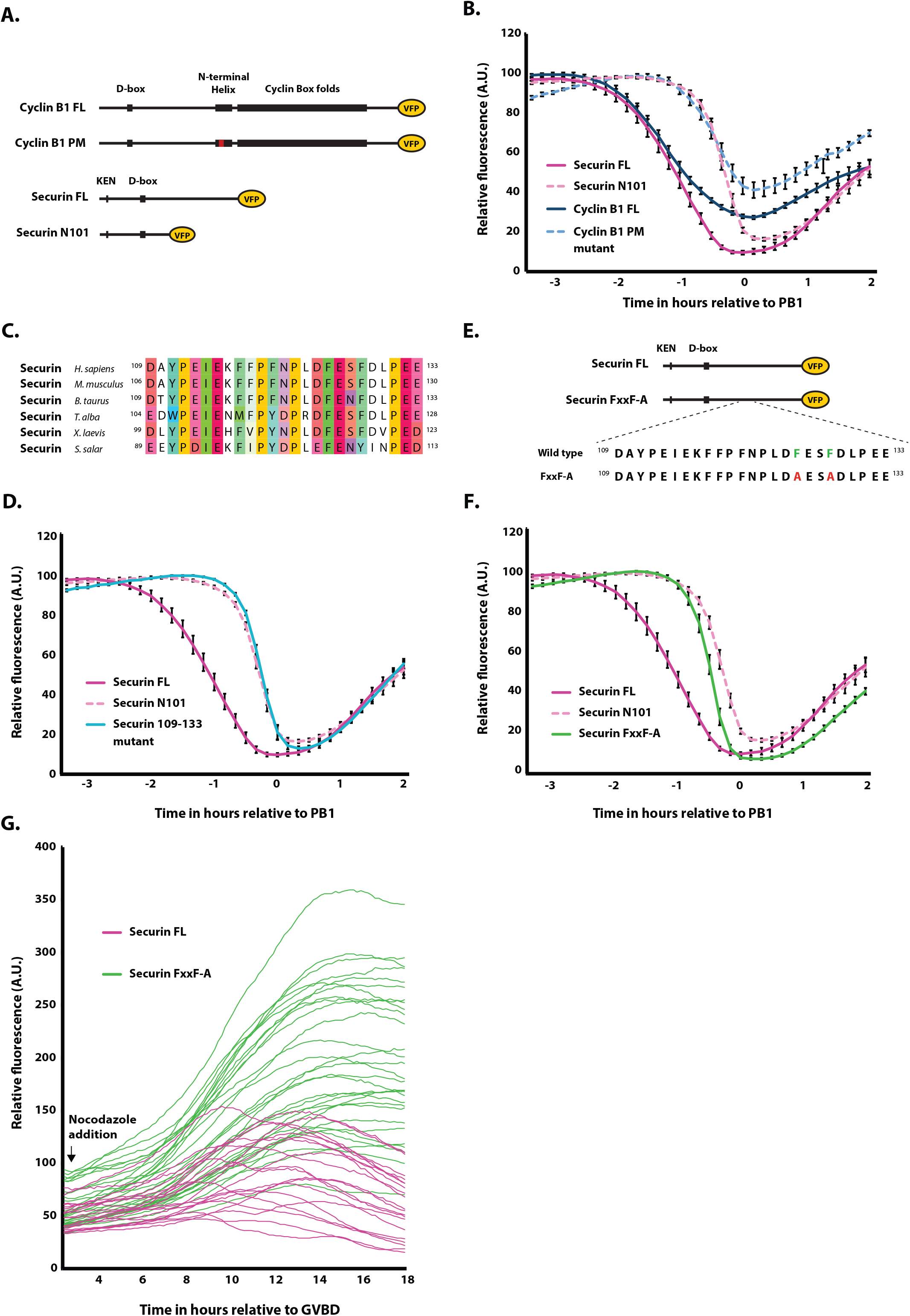
A discrete region within the C-terminus of securin promotes destruction in prometaphase. (A) Schematic showing VFP-tagged securin and cyclin B1 truncations and mutations. (B) Average securin FL (magenta trace, n = 25), securin N101 (pink dashed trace, n = 23), cyclin B1 FL (blue trace, n = 32) and cyclin B1 PM mutant (light blue dashed trace, n=34) destruction profiles relative to PB1 extrusion. (C) Sequence alignment of residues 109-133 in securin orthologs. (D) Average VFP-tagged securin FL (magenta trace, n = 25), securin N101 (pink dashed trace, n = 23) and securin 109-133 mutant (light blue trace, n = 23) destruction profiles relative to PB1 extrusion. (E) Schematic showing the position of Securin FxxF-A amino acid substitutions. Residues F125 and F128 (shown in green in the wild type protein) were switched to alanines (shown in red in Securin FxxF-A). (F) Average VFP-tagged securin FL (magenta trace, n = 25), securin N101 (pink dashed trace, n = 23) and securin FxxF-A (green trace, n = 20) destruction traces relative to PB1 extrusion. (G) VFP-tagged securin FL (magenta traces, n = 20) and securin FxxF-A (green traces, n = 30) destruction profiles following incubation in 150 nM nocodazole to arrest oocytes in prometaphase. These oocytes do not extrude a polar body and are therefore aligned at GVBD. Error bars ± SEM.

Strikingly, we found that securin FL was consistently targeted for destruction earlier than securin N101 (~80 minutes; Fig. 2b). Furthermore, when we compared the data generated in our cyclin B1 study, we found that securin N101 destruction was restricted to metaphase in time with the cyclin B1 mutant lacking the PM motif. This observation raised the possibility that, similar to cyclin B1 and other prometaphase APC/C substrates, an additional region is necessary to act alongside the D-box and direct wild-type securin degradation in prometaphase I in oocytes.

### A discrete region within the C-terminus of securin promotes destruction in prometaphase

To test if a second destruction motif exists within securin to facilitate prometaphase destruction, we replaced a highly conserved C-terminal region located between residues 109-133 (Fig. 2c) with a neutral 25 amino acid TGSGP repeat linker in the full-length securin construct (securin 109-133 mutant). This construct was targeted for destruction in time with securin N101 (Fig. 2d), ~80 minutes after securin FL. This indicated that residues between 109-133 are essential for prometaphase securin destruction in oocyte MI.

Following this finding, we divided residues 109-133 into three groups based on sequence conservation, mutating each group to assess their importance in the timing of securin degradation (Supplementary fig. 1a). Based on sequence similarity with the PM motif in cyclin B1, which centres on residues ^170^DIY^172^ in the mouse ortholog, we predicted that the first mutant, securin DAYPEIE-A, would eliminate prometaphase targeting. Surprisingly however, securin DAYPEIE-A was instead targeted in time with securin FL (Supplementary fig.1b). In contrast, both FFPFNP-A and DFESFD-A mutations resulted in dramatic shifts in destruction timing (Supplementary fig.1c-d). Securin FFPFNP-A was targeted for destruction approximately 60 minutes after securin FL but still ~20 minutes ahead of securin N101 (Supplementary fig.1c), while securin DFESFD-A destruction mirrored that of securin N101 (Supplementary fig. 1d). This suggests that residues essential for wild-type prometaphase securin degradation lie within both of these regions. Since securin DFESFD-A showed the most striking phenotype, we mutated a pair of highly conserved phenylalanines (F125 and F128) to alanines (securin FxxF-A; Fig. 2e). Here, degradation was delayed by 90 minutes, mimicking securin N101 (Fig. 2f; see supplementary fig. 1e for destruction timings of all mutants). Importantly, mutation of residues F125 and F128 to alanines did not impair securin binding to separase as shown by peptide pull-down (Supplementary fig. 1f). We suggest that these two phenylalanine residues are a crucial component of a novel interaction that mediates the timing of securin loss by permitting prometaphase destruction in MI oocytes. In the absence of this interaction, securin destruction initiates 90 minutes later and only in metaphase. Interestingly, these two residues and their surrounding region have no sequence similarity with the PM motif present in cyclin B1.

To confirm that the difference in degradation timing between securin FL and securin FxxF-A was not simply due to differences in protein expression, oocytes were treated with cycloheximide (CHX) to block protein synthesis (Supplementary fig. 2). Upon addition of CHX, the securin FL protein turnover was evident. Approximately 20% of the total protein was lost before a steep period of destruction began 4 hours after CHX addition. In contrast, securin FxxF-A levels were relatively stable until destruction began ~5.5 hours after CHX addition. Critically, securin FxxF-A destruction begins around 90 minutes after securin FL, consistent with results from non CHX-treated oocytes.

Securin FL destruction in MI mouse oocytes begins in prometaphase at a time when Mad2 staining is still detectable at kinetochores^40^. This suggests that the SAC might have less influence over securin FL than securin FxxF-A. To assess this, we treated oocytes with 150 nM nocodazole. This dose of nocodazole depolymerises microtubules and activates the SAC such that PB1 extrusion is blocked in >98% of oocytes. Under these conditions, the rate of securin FL degradation was dramatically reduced, though it was still almost fully degraded within a 10-hour period. In contrast, securin FxxF-A was almost completely stabilised (Fig. 2g). Together our data suggest that an interaction involving two key phenylalanine residues within securin is able to bypass a SAC signal capable of blocking destruction of substrates where only the D-box is accessible. The D-box is however essential for both phases of securin destruction, since mutation of the D-box perturbs both metaphase and prometaphase securin destruction, and inhibits PB1 extrusion (Supplementary fig. 3c).

Our data are consistent with a biphasic securin destruction process in MI oocytes. A first period of destruction 3 hours ahead of PB1 extrusion which requires both the D-box and an additional region with an absolute requirement for residues F125 and F128. This is followed by a second period of destruction in metaphase as the D-box becomes sufficient for degradation 1 hour ahead of PB1 extrusion. Critically, our separase biosensor data indicates that separase is only active during the latter half of the second phase of securin destruction.

### Two phenylalanine residues key to prometaphase securin destruction are predicted to be masked when securin is bound to separase

Interestingly, F125 and F128 are positioned within securin’s separase interaction segment, in the vicinity of the residues involved in securin’s pseudosubstrate binding to the separase active site^41,42^. We therefore asked how these phenylalanine residues are positioned when securin is in complex with separase. Since a mammalian crystal structure for the securin-separase complex is yet to be solved, we instead used the existing structure of the *S. cerevisiae* complex as a reference^41^. Critically, these residues and the surrounding region are largely conserved between mammals and yeast (Fig. 3a). In the structure, residues Y276 and F279, corresponding to F125 and F128 in the human protein, sit deep within a hydrophobic binding pocket on the surface of separase (Fig. 3b). It is therefore highly likely that these two residues would be obscured when securin is in complex with separase. The conservation of these residues throughout eukaryotes (Fig. 3a), coupled with their neighbouring position to the pseudosubstrate region of securin, lead us to propose that these residues would be similarly hidden in other species.

**Figure 3.**
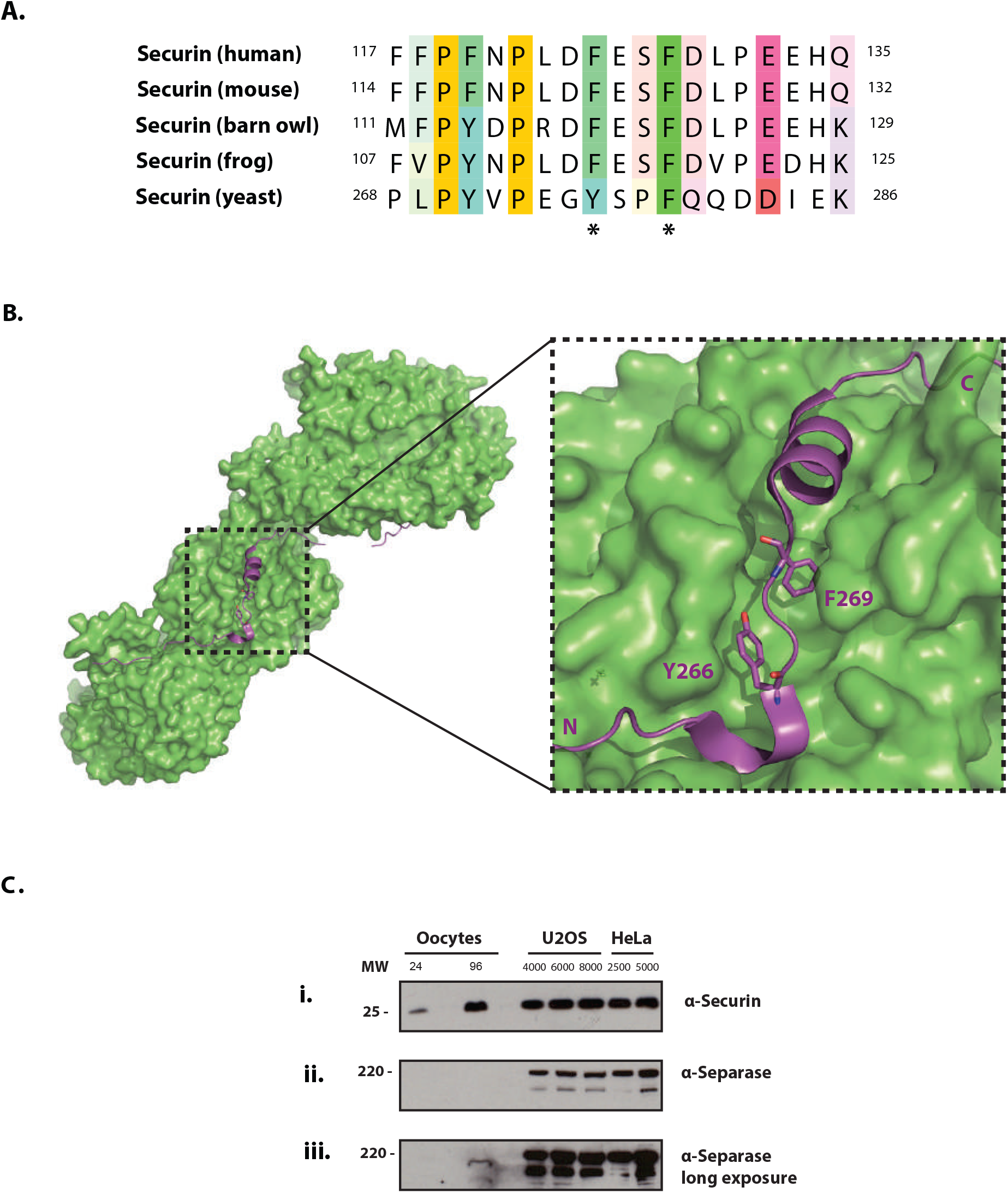
F125 and F128 are predicted to direct the prometaphase destruction of a pool of non-separase-bound excess securin. (A) Alignment of the region surrounding residues F125 and F128 in securin orthologs. (B) The molecular surface of separase (green) in contact with the region of securin detailed in Fig. 3a (purple). Image generated using the crystal structure of the *Saccharomyces cerevisiae* separase-securin complex. The side chains of securin residues Y276 and F279, which correspond to F125 and F128 in the human protein, are shown as stick models and labelled in purple. The N- and C-termini of the securin segment displayed are also labelled in purple (C) Western blot analysis of lysates prepared from mitotic cells in late prometaphase (U2OS and HeLa cells), and mouse oocytes in late prometaphase (collected at 5.5 hours post GVBD). The numbers of mitotic cells and oocytes loaded into each lane are as indicated. Membranes were used to detect (i) securin, (ii) separase by a short exposure and (iii) separase by a long exposure (iii).

### Securin is present in large excess over separase in mouse oocytes

Given that F125 and F128 are predicted to be masked when securin is bound to separase, we speculated that destruction in prometaphase must represent that of a non-separase bound population of securin. In mitosis, securin protein is in excess of separase^43,44^. This ratio has been quantified in HeLa cells, where free securin is reported to be 4-5 fold more abundant than separase-associated securin^43^. Therefore we considered that a similar excess of securin in oocytes could provide the basis for a mechanism able to support an extended period of securin destruction without a release of separase inhibition. To address this, we compared the separase to securin ratio in mitosis with that in MI oocytes by western blotting. We find that securin is present at comparable levels between 96 oocytes and HeLa cell extracts prepared from 2500 and 5000 cells (Fig. 3ci). Yet while separase was readily detectable in the mitotic extracts, no band was observed in the 96 oocyte lane (Fig. 3cii). By extending the exposure time from 30 seconds to 10 minutes, a small separase band was detected in the 96 oocyte lane, where all mitotic lanes became strongly overexposed (Fig. 3ciii). We conclude that in oocytes, securin is present in much greater excess to separase than the 4-5 fold difference documented in HeLa cells.

We therefore suggest a mechanism by which a large pool of excess non-separase bound securin is preferentially targeted for destruction in late prometaphase by a mechanism involving both the D-box and a second region where F125 and F128 are essential. Following this, a smaller pool of separase-bound securin becomes a destruction target in metaphase.

Preferential destruction of non-separase bound securin has previously been demonstrated in HeLa cells. Hellmuth et al. showed that separase-bound securin is dephosphorylated by PP2A-B56 phosphatase, whereas free securin exists in a phosphorylated state and is a preferential APC/C target^43^. Importantly however, the destruction of both separase-bound and non-separase bound securin in mitosis takes place only after SAC satisfaction. It therefore seems unlikely that this same mechanism is responsible for degradation of non-separase bound securin in prometaphase in mouse oocytes. To confirm this, we mutated four key phosphorylation sites in securin to alanines (securin 4A; S31A, T66A, S87A and S89A) and observed no change in destruction timing compared to securin FL (Supplementary fig. 4). We therefore suggest that the mechanism of non-separase bound securin degradation described in this manuscript occurs independently of phosphorylation status.

### Two complimentary mechanisms prevent premature separase activation and division errors

We next wanted to investigate how separase activity is affected in meiosis when the large excess of securin is removed. Chiang et al. previously showed that segregation errors in fixed oocytes were only observed when both securin and cyclin B1-Cdk1-mediated separase inhibition were removed^19^. However, when these errors arise and how separase is regulated through time remains unclear. To investigate this, we knocked down securin protein expression with a morpholino oligo (MO) such that in prometaphase, MO oocytes contained ~13% of the protein level relative to non-treated control cells (quantified by western blot; Supplementary fig. 5). Despite severe depletion of securin, the cleavage activity of separase was still restricted to the final 30 minutes preceding PB1 extrusion (Fig 4a). This result agrees with observations made in fixed oocytes, suggesting that cyclin B1-Cdk1 levels are sufficient to compensate for the loss of securin. To test whether this was also the case in our live cell assay, we injected oocytes with a previously published separase AA mutant^19^. This construct is fully active but lacks Cdk1 phosphorylation sites (S1121A and T1342A) that prevent its inhibition by cyclin B1-Cdk1. Importantly, oocytes expressing separase AA protein produce separase activity profiles identical to control oocytes, suggesting that securin was similarly sufficient to compensate for a loss of cyclin B1-Cdk1-mediated inhibition (Fig. 4a). In contrast, where securin protein levels are restricted in separase AA expressing oocytes (separase AA + securin MO), cleavage activity initiates up to 1 hour ahead of polar body extrusion (Fig. 4a). Furthermore, by the time they extruded a polar body, separase AA + securin MO-injected oocytes had cleaved roughly twice as much sensor compared to control cells (Fig. 4a).

**Figure 4.**
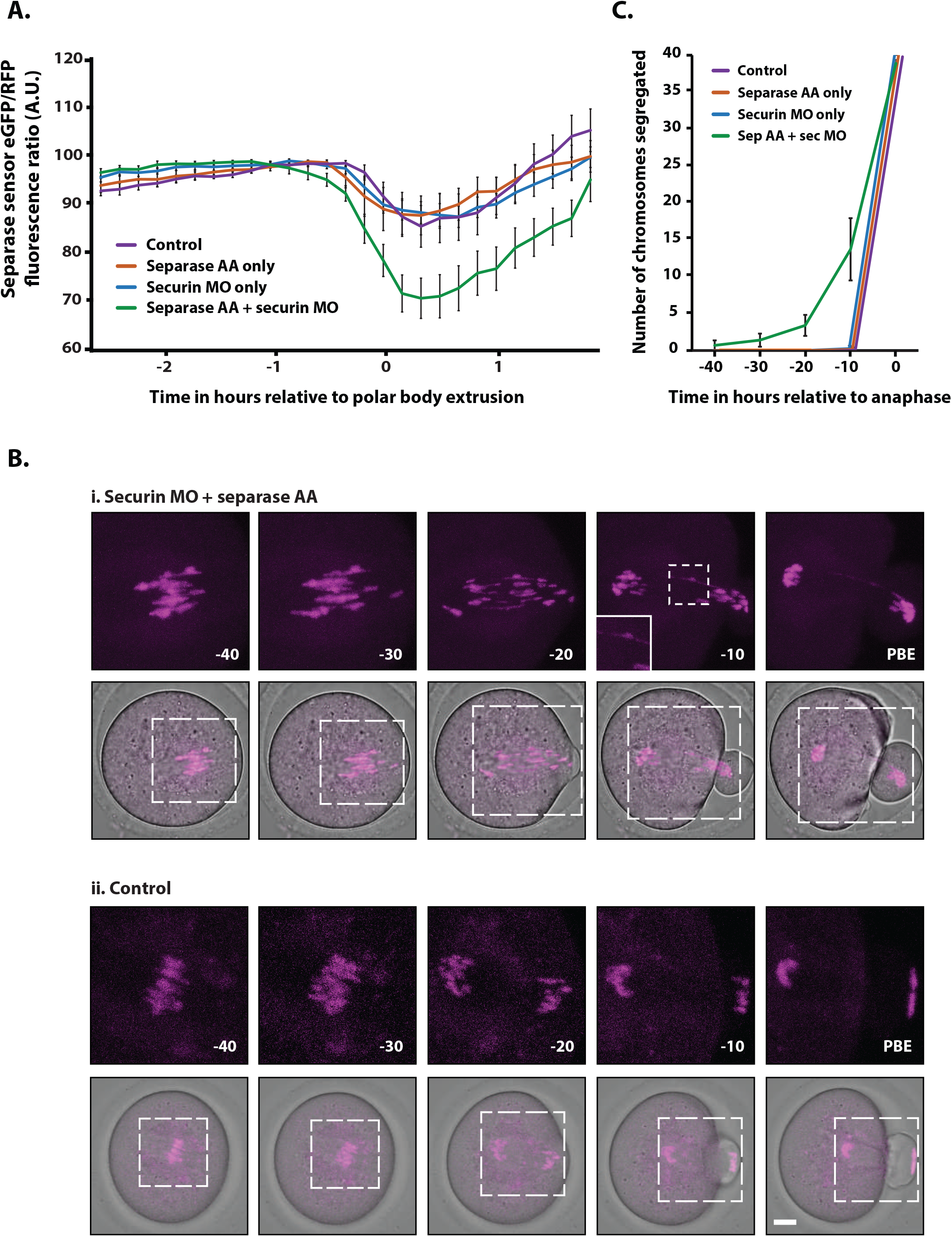
Both a securin excess and cyclin B1-Cdk1-mediated inhibition are independently sufficient to suppress separase activity and prevent segregation errors in mouse oocyte meiosis I. (A) Graph showing mean MI separase activity profiles as determined by a separase activity biosensor (eGFP/RFP ratio) for control (purple trace, n = 8), separase AA-injected (a fully active separase construct lacking Cdk1 phosphorylation sites S1121A and T1342A; orange trace, n = 7), securin MO-injected (blue trace, n = 8) and separase AA + securin MO-injected (green trace, n = 10) mouse oocytes relative to PB1 extrusion. Error bars ± SEM. (B) Representative images showing chromosome segregation in the 40 minutes prior to PB1 extrusion in (i) securin MO + separase AA-injected and, (ii) control oocytes. Chromosomes were visualised by incubating oocytes with SiR-DNA (shown in magenta and merged with bright field) at times shown relative to PB1 extrusion. Scale bar = 10µm. SiR-DNA images correspond to the areas marked by the white dotted lines in the merged images. (C) Graph showing the mean number of chromosomes segregated per 10 minute time point in the 40 minutes prior to anaphase (defined as the time point in which more than 50% of chromosomes had segregated) for control (purple trace, n = 8), separase AA-injected (orange trace, n = 7), securin MO-injected (blue trace, n = 8) and separase AA + securin MO-injected (green trace, n = 10). Error bars ± SEM.

To monitor the impact of premature separase activity in separase AA + securin MO oocytes, we used confocal microscopy to image live oocytes stained with SiR-DNA to track individual chromosome movements^45^. This analysis revealed premature chromosome segregation from 40 minutes ahead of polar body extrusion in separase AA + securin MO oocytes resulting in a non-synchronous anaphase and an increased occurrence of lagging chromosomes (Fig 4bi + supplementary videos 1-4). Furthermore, in 13% of separase AA + securin MO oocytes, the segregation errors were so severe that anaphase failed completely (supplementary videos 5 and 6). This is in stark contrast to the tightly synchronised anaphase observed in untreated control oocytes ~20 minutes ahead of polar body extrusion (Fig 4bii + supplementary videos 7-9). Quantification of the number of chromosomes segregated in each 10-minute time interval prior to anaphase (where anaphase was defined as the time point at which more than 50% of chromosomes had segregated) revealed that the premature separation of bivalents only took place in separase AA expressing oocytes where securin levels were restricted. In either separase AA only, or securin MO only treatment groups, almost all oocytes segregated their chromosomes synchronously within a single 10-minute time interval (Fig 4c).

These data further demonstrate that either inhibitory pathway, an excess of securin or inhibition by cyclin B1-Cdk1, is sufficient to prevent premature separase activity in mouse oocytes. When both pathways are diminished, the timing of anaphase and the timing of separase release are uncoupled. This results in severe chromosome segregation defects. Interestingly, even when both a securin excess is depleted and cyclin B1-Cdk1-mediated inhibition is prevented, separase activity still remains suppressed until the final hour of meiosis. This suggests that even severely restricted securin levels are sufficient for separase inhibition during the hours leading up to metaphase in meiosis.

## Discussion

Similar to sister chromatid segregation in mitosis, the segregation of homologous chromosome pairs in mouse oocyte meiosis I requires the proteolysis of both securin and cyclin B1^23^. Importantly, in both somatic and germ cells, the degradation of securin and cyclin B1 is synchronous and both proteins must be depleted in order for separase activation and Cdk1 inactivation to be temporally coupled^23–25^. In mouse meiosis I, a premature loss of Cdk1 activity ahead of securin degradation drives polar body extrusion, yet chromosomes do not segregate and instead become trapped within the cleavage furrow^46^.

Here we show that mouse oocytes in meiosis I contain a large excess of securin over separase (several fold beyond that reported in mitosis). We reveal the existence of a novel mechanism of targeted degradation that functions to promote the destruction of non-separase bound securin in prometaphase. Critically this mechanism relies on a key region of securin which is only exposed when securin is not bound to separase. We suggest that in oocytes, the majority of non-separase-bound securin is removed by this mechanism, while a much smaller fraction of inhibitory separase-bound securin is only targeted in metaphase. By this strategy, the cellular destruction machinery is far less likely to be overwhelmed by excessive securin molecules during the final stages of chromosome segregation. This is particularly important given that separase must be rapidly released simultaneously on all chromosomes, ensuring their synchronous segregation. Indeed, our recent data demonstrates a similar strategy for the removal of non-Cdk1-bound cyclin B1 ahead of cyclin B1-Cdk1 activity; essential to permit a rapid drop in Cdk1 activity and to prevent anaphase from stalling^36^.

We show that securin destruction is biphasic in MI oocytes, consisting of a metaphase period of D-box only destruction (resembling mitotic destruction), and a prometaphase period of destruction requiring both the D-box and an additional region able to bypass spindle checkpoint inhibition. How this additional region functions to bypass the spindle checkpoint remains unclear. However we suggest that, similar to other prometaphase APC/C substrates, this region of securin could be involved in either direct binding to the APC/C or in outcompeting spindle assembly checkpoint protein inhibition of APC/C-Cdc20^47–50^. In addition, cyclin B3 has recently been shown to have an important role in mouse oocyte MI in priming the APC/C for securin and cyclin B1 destruction^39,51,52^. Therefore an alternative possibility is that this region in securin is regulated by cyclin B3 and involved in an oocyte-specific APC/C activation mechanism. It is worth noting that the regions we describe in securin and cyclin B1 do not share sequence similarity. It is therefore possible that while their destruction timings are coupled, the mechanisms by which they are targeted early in prometaphase may be distinct.

Chiang et al. report in fixed oocytes that either securin or cyclin B1-Cdk1 inhibitory pathways are sufficient to independently prevent chromosome segregation errors in young mouse oocytes^19^. Our results in live oocytes agree with this, and further demonstrate that separase activity is only released prematurely when both inhibitory pathways are perturbed. It remains to be determined whether the excess of non-separase-bound securin becomes essential in aged mouse oocytes (where cohesin levels are significantly reduced^53,54^), and over the prolonged duration of prometaphase in human oocytes (14 – 18 hours compared to 4.5 - 6.5 hours in mouse^55,56^). Furthermore, it seems likely than an excess of securin in MI, could function to ensure a threshold level of securin protein in MII. MII securin protein level is particularly important since unlike MI, cyclin B1-Cdk1 is no longer able to compensate for depleted securin in this division^13,19^. Indeed Nabti et al. revealed that oocytes from aged mice destroy securin more rapidly and to a greater extent than oocytes from younger mice. As a consequence, separase inhibition in MII is incomplete in these cells and sister chromatid cohesion is lost prematurely resulting in segregation defects^57^.

Destruction of securin during prometaphase has previously been considered as precocious, an error resulting from a weakened SAC in oocytes^58^. Our work instead demonstrates that this period of destruction is deliberate. In prometaphase, an excess of non-separase bound securin is targeted for destruction through a novel mechanism in oocytes. By this strategy, separase bound securin is protected, allowing for a switch-like activation of separase, essential for the fidelity of anaphase in oocytes.

## Supporting information

Supplementary video 1 - separase AA + securin MO - Lagging chromosomes + chromosome bridge

Supplementary video 2 - separase AA + securin MO - Premature segregation

Supplementary video 3 - separase AA + securin MO - Premature segregation + lagging chromosomes

Supplementary video 4 - separase AA + securin MO - Premature segregation

Supplementary video 5 - separase AA + securin MO - Anaphase failure

Supplementary video 6 - separase AA + securin MO - Anaphase failure

Supplementary video 7 - securin MO only

Supplementary video 8 - separase AA only

Supplementary video 9 - Control

---

**Supplementary figure 1.**
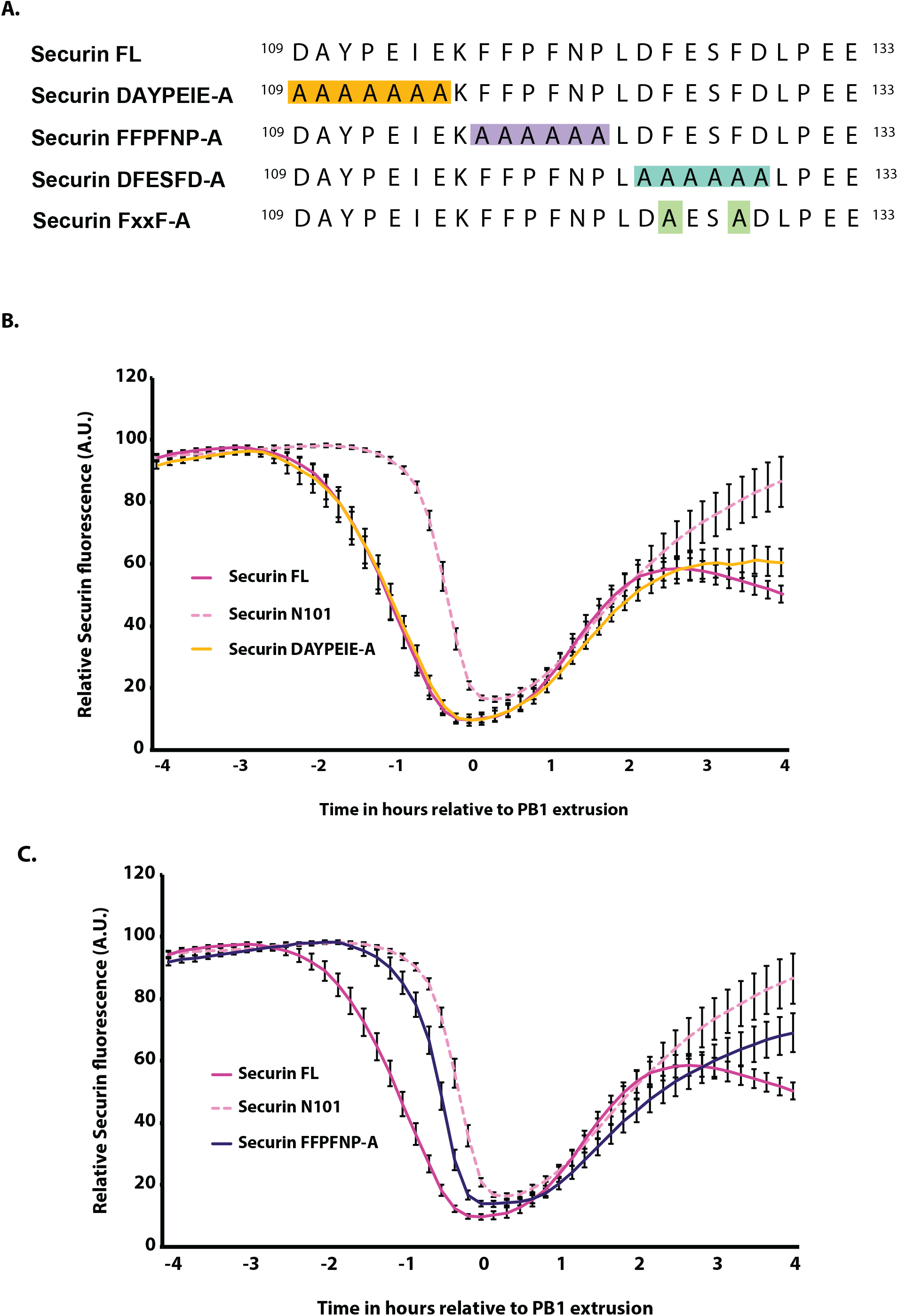
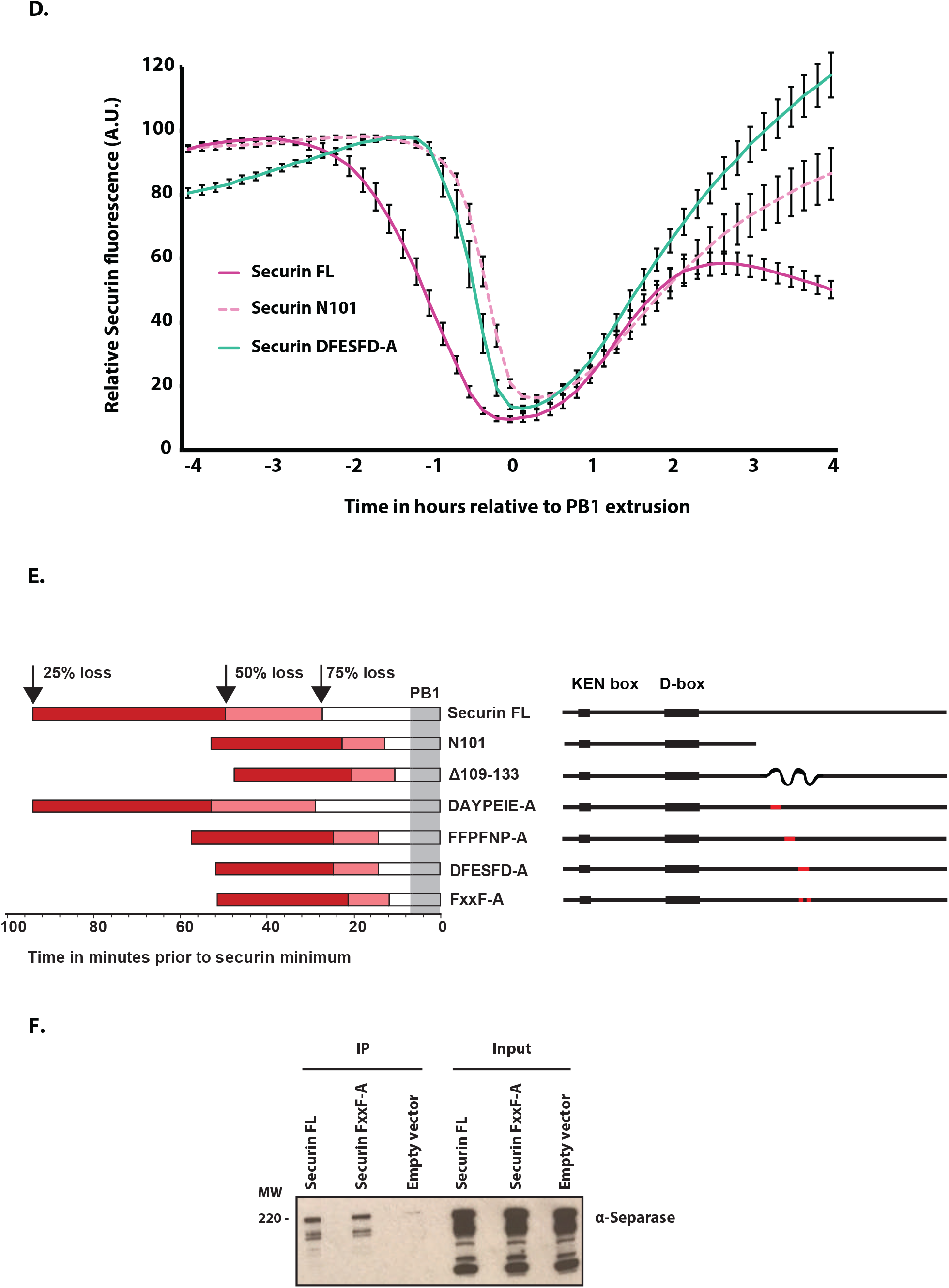
Destruction profiles of securin mutation constructs leading to identification of the F125 and F128 as essential for prometaphase securin destruction. (A) Securin residues 109-133 showing sequence detail relative to the nomenclature of all venus-tagged securin mutations assayed in parts B-D. (B) Average VFP-tagged securin FL (magenta trace, n = 25), securin N101 (dashed pink trace, n = 23) and securin DAYPEIE-A (yellow trace, n = 19) destruction traces. (C) Average VFP tagged securin FL (magenta trace, n = 25), securin N101 (dashed pick trace, n = 23) and securin FFPFNP-A (purple trace, n = 23) destruction traces. (D) Average VFP-tagged securin FL (magenta trace, n = 25), securin N101 (dashed pick trace, n = 23) and securin DFESFD-A (green trace, n = 20) destruction traces. All traces are lined at PB1 extrusion and error bars = ±SEM throughout. (E) Direct comparison of the timing of destruction of all Venus-tagged securin truncations and mutants plotted in parts B-D. Schematic representations of securin constructs are shown down the right hand side. To the left, the length of each bar indicates each construct’s destruction timing relative to complete destruction (time 0; approximately 0-10 mins post PB1 extrusion). The open, white bars indicate the point at which 75% of the destruction has taken place. The light red extension to this bar indicates the point at which 50% of the destruction has taken place, followed by a dark red extension indicating the point at which 25% of the destruction has taken place. The period over which PB1 extrusions occur is shaded in grey. (F) HeLa cells transfected with securin FL, securin FxxF-A or empty mVenus N1 transfection vector as indicated were synchronised in nocodazole for 16 hours and collected by mitotic shake off. Cells were then lysed and anti-GFP immunoprecipitates (IP) were probed for separase. Input signals are also shown.

**Supplementary figure 2.**
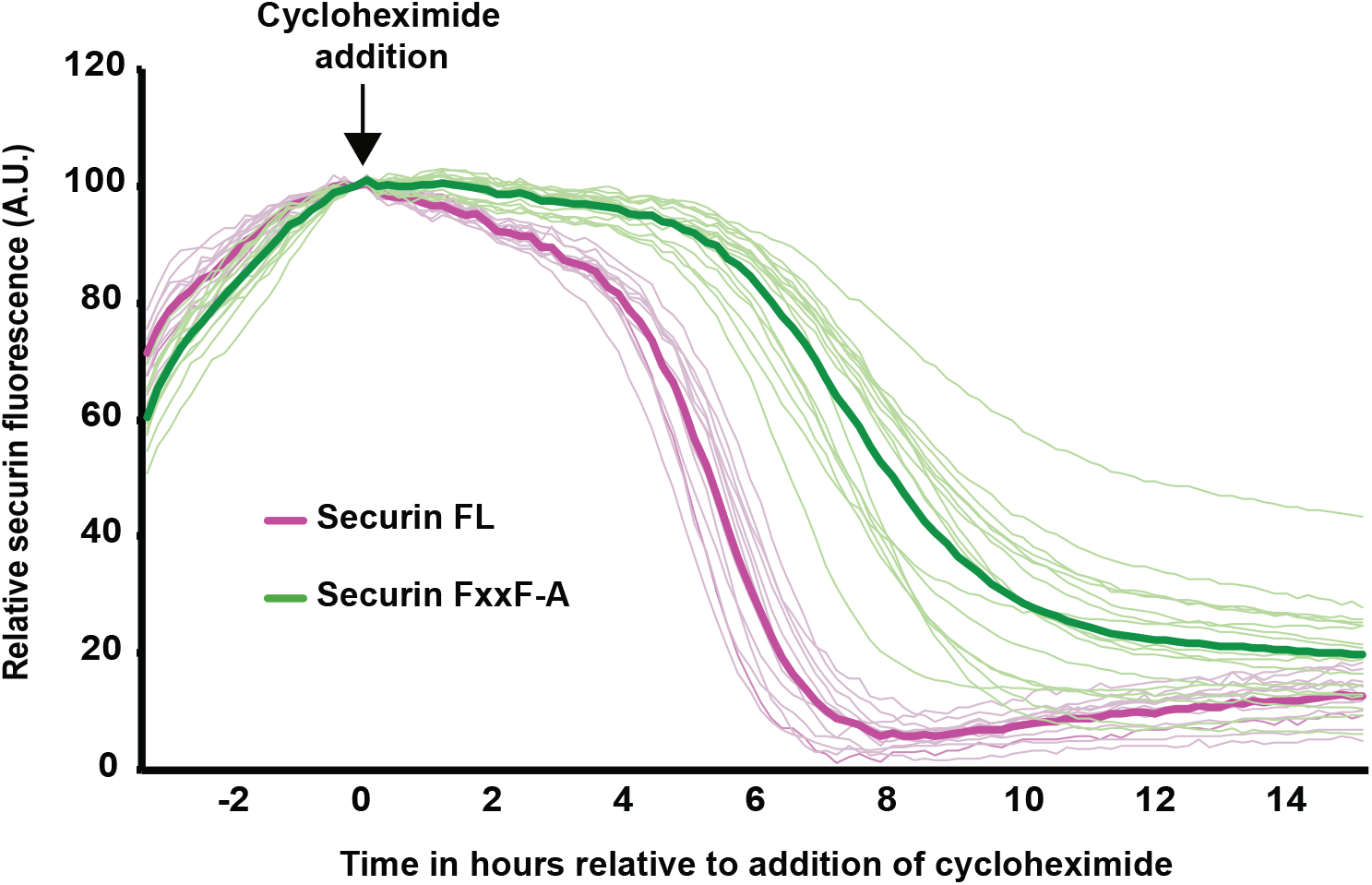
Differences in the timing of securin FL and FxxF-A destruction are not due to a difference in protein expression rate. Average VFP-tagged securin FL (magenta traces, n = 24) and securin FxxF-A (green traces, n = 27) on addition of cycloheximide to inhibit protein synthesis. Traces are aligned to the addition of cycloheximide at 3 hours post GVBD. Fine traces represent destruction profiles from individual oocytes; heavy traces represent the average destruction profile resulting from all injected oocytes of a given construct.

**Supplementary figure 3.**
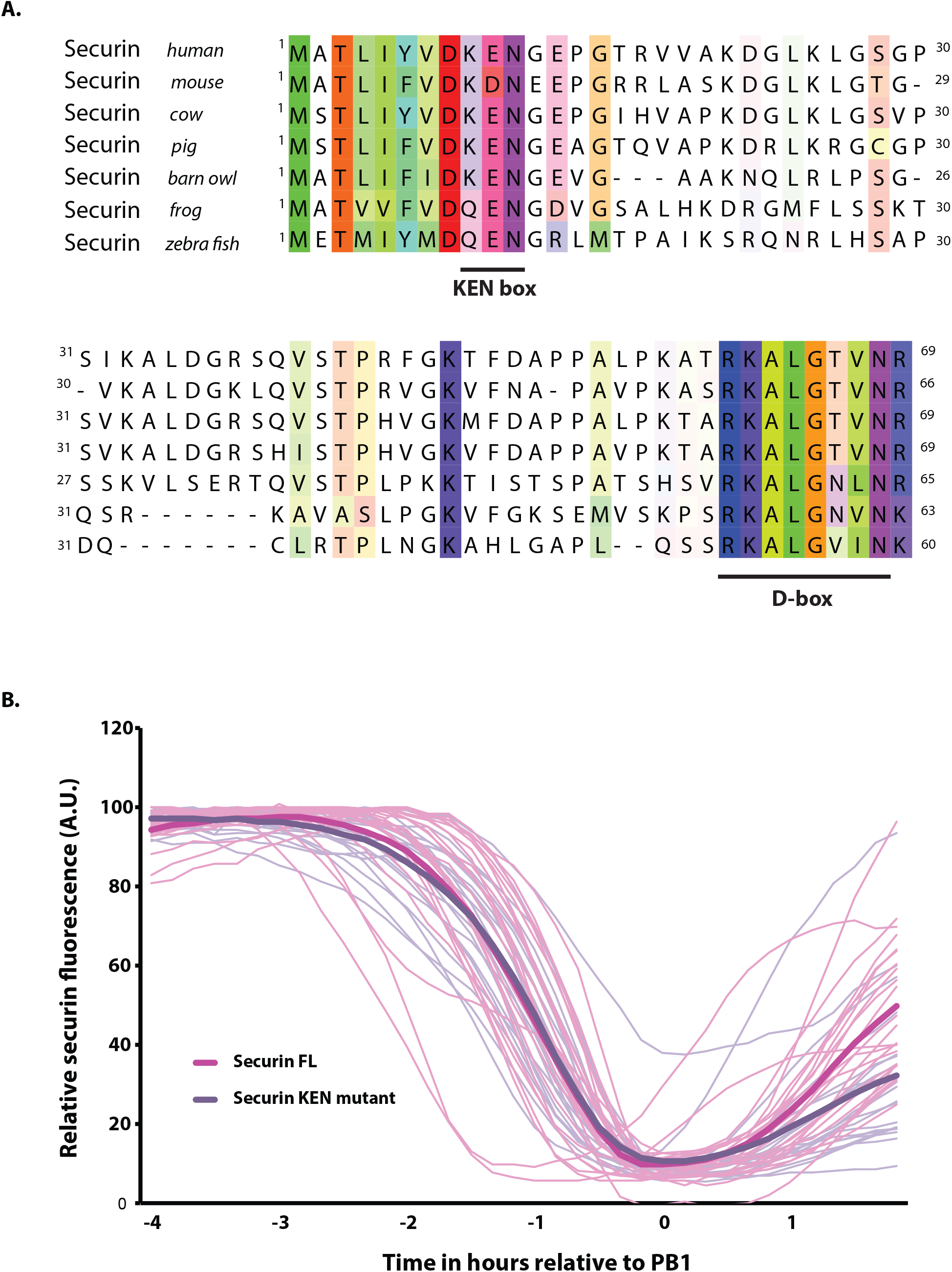
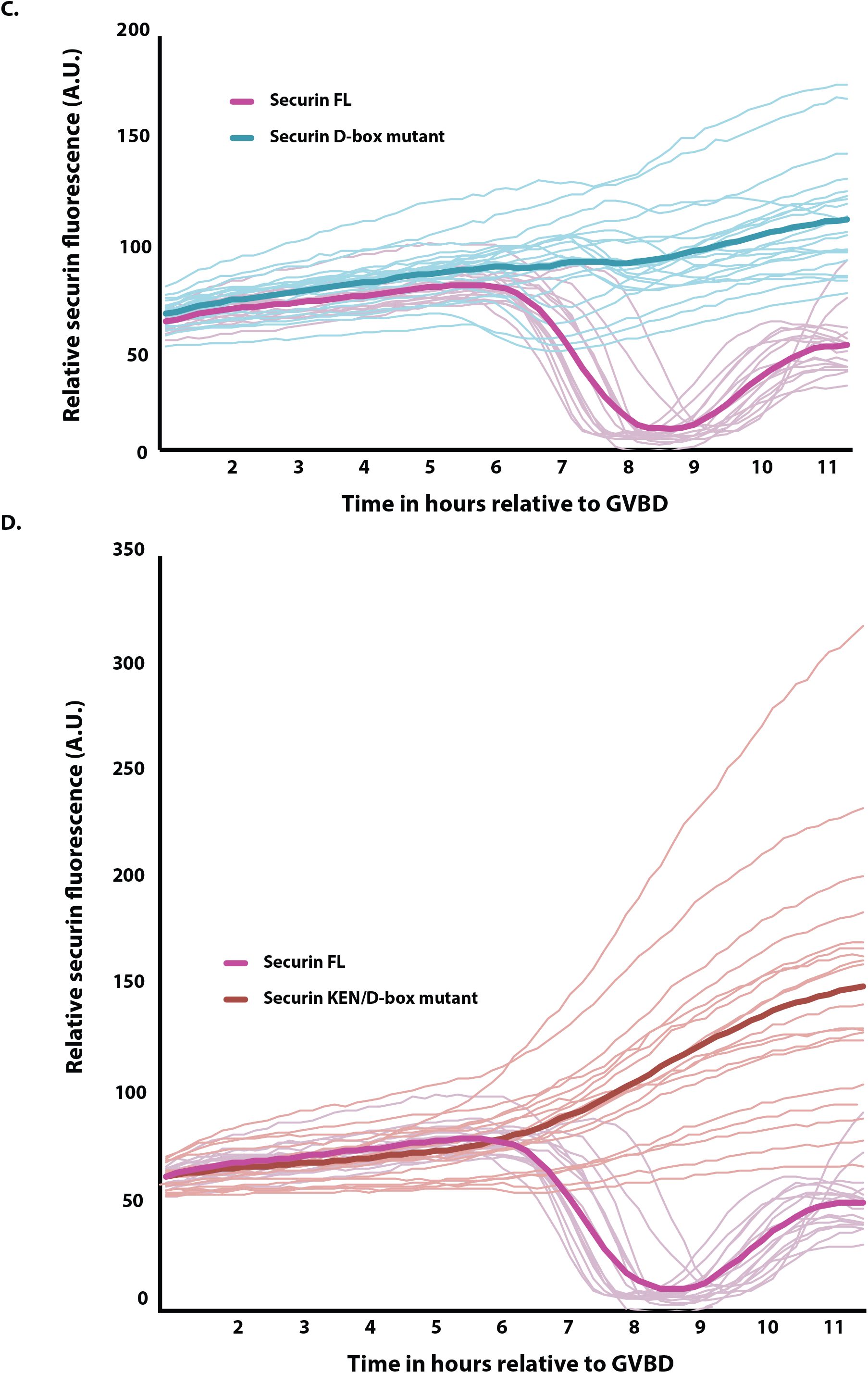
Meiotic securin destruction is D-box dependent but not KEN box dependent. (A) Alignment of residues 1-69 in securin orthologs containing both KEN box and D-box motifs as indicated. (B) Average VFP-tagged securin FL (magenta traces, n = 25) and securin KEN mutant (purple traces, n = 20) destruction profiles aligned at PB1 extrusion. (C) Average VFP-tagged securin FL (magenta traces, n = 16) and securin D-box mutant (light blue traces, n = 23) destruction profiles aligned at GVBD (oocytes expressing D-box mutant securin arrest their cell cycle in metaphase do not extrude a polar body). (D) Average VFP-tagged securin FL (magenta traces, n = 16) and securin KEN/D-box mutant (red traces, n = 19) destruction profiles aligned at GVBD. Note that though neither securin D-box mutant nor securin KEN/D-box mutant expressing oocytes extrude a polar body, securin destruction is only completely inhibited where both the D- and the KEN-box are absent. Fine traces represent individual oocyte fluorescence levels and heavy traces represent their average throughout.

**Supplementary figure 4.**
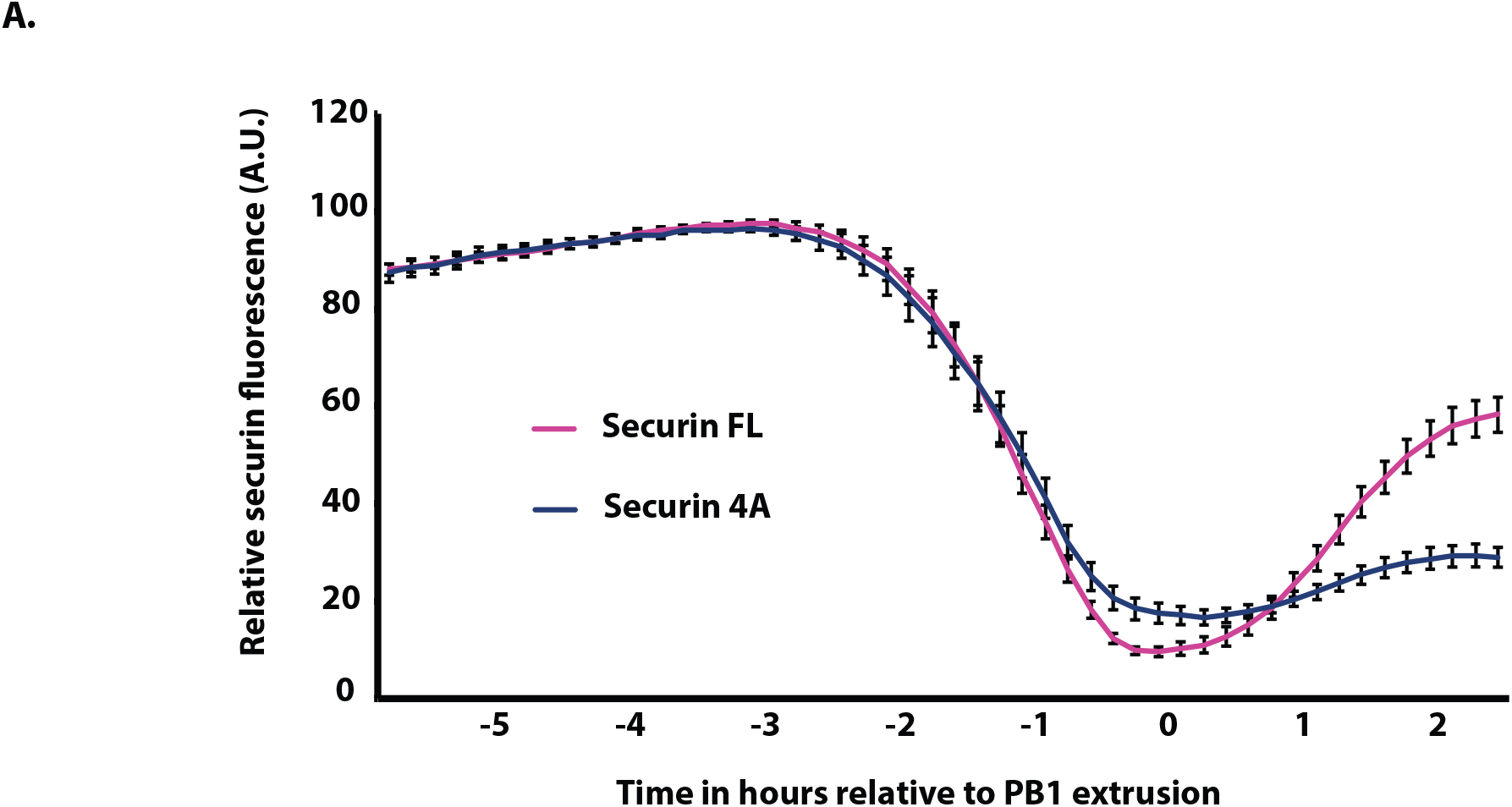
A securin phosphomutant does not affect degradation timing in meiosis I mouse oocytes. (A) Average VFP-tagged securin FL (magenta trace, n=25) and securin 4A (blue trace, n=17; securin 4A is a phosphomutant containing mutations S31A, T66A, S87A and S89A^43^) destruction profiles aligned at PB1 extrusion. Error bars = +/− SEM.

**Supplementary figure 5.**
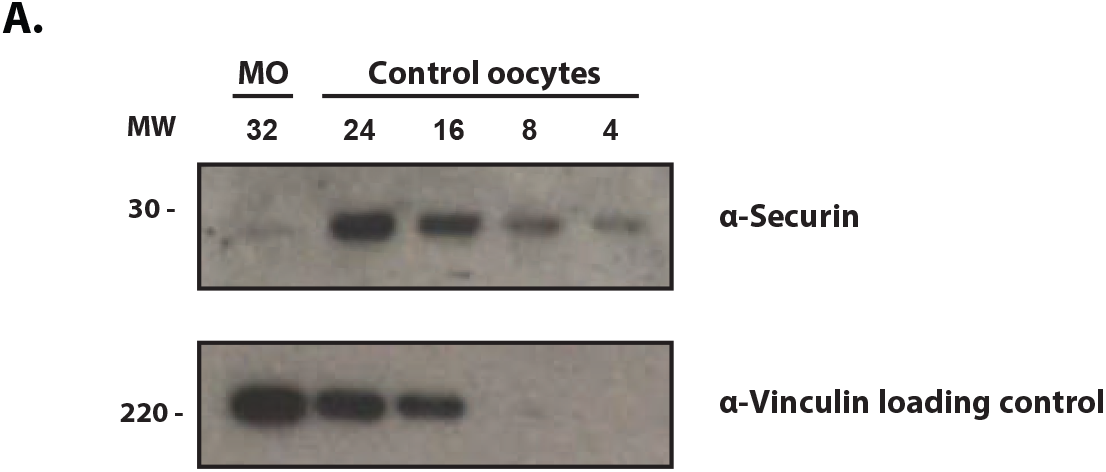
Quantification of securin morpholino knock down. Western blot of control and securin morpholino (MO) injected oocytes collected 5.5 hours post GVBD (numbers of oocytes loaded per lane are indicated). Quantification of protein bands indicates an 87% knockdown of securin in MO-injected oocytes. The lower blot was probed for vinculin and used as a loading control.

## Methods

### Contact for Reagent and Resource Sharing

Further information and requests for resources and reagents should be directed to Suzanne Madgwick.

### Gamete Collection and Culture

4 to 8-week-old female, outbred, CD1 mice (Charles River) were used. All animals were handled in accordance with ethics approved by the UK Home Office Animals Scientific Procedures Act 1986. GV stage oocytes were collected from ovaries punctured with a sterile needle and stripped of their cumulus cells mechanically using a pipette. For bench handling, microinjections and imaging experiments, oocytes were cultured at 37°C in medium M2 (Sigma) supplemented where necessary with the addition of 30 nM 3-isobutyl-1-methylxanthine (IBMX; Sigma) to arrest oocytes at prophase I. Data was only collected from oocytes that underwent GVBD with normal timings and had a diameter within 95-105% of the population average. To ensure reproducibility, oocyte data sets were gathered from a minimum of 3 independent experiments. For each independent experiment, both control and treatments groups were derived from the same pool of oocytes, collected from a minimum of 2 animals. Oocytes were selected at random for microinjection. Where necessary, and at the times indicated, nocodazole (Sigma) was added to the media at a final concentration of 150 nM. For confocal imaging SiR Hoechst was added to media 30 minutes prior to imaging at a final concentration of 250 nM^45^.

### Preparation of cRNA constructs for microinjection

Microtubule associated protein 7 (Map7; to visualise microtubules^59^), wild type human securin, securin truncations, separase biosensor (a gift from Jan Van Deursen) and separase AA mutant (a gift from Michael Lampson) sequences were amplified by PCR as previously described^60^. Mutations within securin were generated by primer overhang extension PCR. Following this we performed Sequence and Ligation Independent Cloning (SLIC)^61^ using modified pRN3 vectors to generate either a construct with no further tags (the separase biosensor) or constructs coupled to a Venus fluorescent protein (all other sequences)^62^. Resultant plasmids were linearized and cRNA for microinjection was prepared using a T3 mMESSAGE mMACHINE kit according to the manufactures instructions (Ambion Inc.). Maximal stability was conferred on all cRNA constructs by the presence of a 5′ globin UTR upstream and both 3’UTR and poly (A)–encoding tracts downstream of the gene. cRNA was dissolved in nuclease-free water to the required micropipette concentration.

### Knockdown of gene expression using morpholinos

A Morpholino antisense oligo designed to recognize the 5’ UTR of securin (sequence: GATAAGAGTAGCCATTCTGGATTAC; MO; Gene Tools) was used to knockdown gene expression. As per the manufacturer’s instructions, the oligo was stored at room temperature, heated for 5 minutes at 65°C prior to use and loaded at a micropipette concentration of 1 mM.

### Microinjection and imaging

Oocyte microinjection of the MO and construct mRNAs was carried out on the heated stage of an inverted microscope fitted for epifluorescence (Olympus; 1×71). In brief, fabricated micropipettes were inserted into cells using the negative capacitance overcompensation facility on an electrophysiological amplifier (World Precision Instruments). This procedure ensures a high rate of survival (>95%). The final volume of injection was estimated by the diameter of displaced ooplasm and was typically between 0.1-0.3% of total volume.

To generate destruction profiles, bright field and fluorescence images were captured every 10 minutes throughout meiosis I by an inverted Olympus IX71 microscope (fitted for epifluorescence) and CCD camera (Micromax, Sony Interline chip, Princeton Instruments). Images were then analysed and processed using MetaFluor software (version 7.7.0.0; Molecular Devices). All experiments were performed at 37°C.

Confocal images (including all experiments using the separase biosensor) were captured using a Zeiss LSM-800. Oocytes were imaged at 10 minute intervals though 20+ Z-sections over a 12-hour period from GV stage. All experiments were performed in a temperature-controlled, humidified chamber set at 37°C. Bright-field and fluorescent images were recorded in Zen Blue (Zeiss) and processed in Fiji. By this method, all oocytes extruded polar bodies.

### Molecular structure images and multiple sequence alignments

Molecular structure images were generated using the PyMOL Molecular Graphics System, version 1.3 Schrödinger, LLC. Sequence conservation alignments were made by importing protein sequences from Uniprot and aligning in Jalview, version 15.0. All figures were prepared in Adobe Illustrator CC, version 17.1.0.

### Mitotic Cell Cultures

Standard laboratory U2OS and HeLa cell lines (a gift from Jonathan Higgins) were used to generate lysates for western blotting. Both cell types were cultured in flasks at 37°C, 5% CO2 in DMEM (Lonza) with 10% FBS (Life Technologies) and antibiotics. Once cell coverage reached ~90%, flasks were treated with 100 nM nocodazole for an 8-hour incubation period for western blotting and a 16-hour incubation period for immunoprecipitation. Metaphase cells were then collected by mechanical shake off and lysed.

### Immunoprecipitation

HeLa cells transfected with securin FL, securin FxxF-A or empty mVenus N1 transfection vector were synchronised and collected as described above. Cells were then lysed in 50 mM Tris-HCl, pH 7.8, 150 mM NaCl, 0.5% NP-40 plus protease inhibitor cocktail (Roche) for 30 min on ice and clarified by a 12,000 g spin for 20 min at 4°C. Complexes were immunoprecipitated for 90 minutes at 4°C with GFP-Trap beads (Chromotek). After five washes in lysis buffer, proteins were eluted from beads by incubating for 10 minutes at 95°C in sample buffer. The supernatant was then analysed by immunoblotting.

### Immunoblotting

Mitotic U2OS and HeLa cell lysates were prepared using Laemmli buffer following mechanical shake off of metaphase cells. Oocytes were collected 5.5 hours after GVBD ± 15 min and lysed in Laemmli buffer. SDS-PAGE and immunoblotting were carried out by standard procedures. Immunoblot membrane sections were incubated for 16 hours at 4°C with either anti-securin (Abcam, AB3305) or anti-separase (Abnova, 6H6). Non-fat milk (5%) was used as a blocking solution and anti–mouse IgG (7076P2; Cell Signaling) and ECL Select (RPN2235; GE Healthcare) were used as secondary detection reagents. ECL Select detection reagents were specifically used to produce x-ray signals with broad linear dynamic range. Membranes were exposed to Hyperfilm x-ray film (Amersham Biosciences) and developed using a SRX101 film processor (Konica). Exposure time depended on the strength of the signal. Immunoblots are representative of 2 independent blots.

### Quantification and Statistical Analysis

Real-time destruction profiles were recorded in MetaFluor (Molecular Devices) and data was automatically logged in Excel. By taking an average VFP intensity reading from a defined region of interest around the oocyte, fluorescence intensity was plotted over time and oocyte data sets were aligned at PB1 extrusion unless otherwise stated. Average polar body extrusion timings were identical between experimental groups unless otherwise stated. Fluorescence data values are arbitrary. In order to give the destruction profile of each oocyte equal weighting, when generating an average treatment destruction profile, all individual data sets were normalised prior to calculation (taking the maximum point of fluorescence prior to protein destruction as 100 a.u.). However, like the examples shown in Figures S2 and S3b-d, we also compared all raw traces, confirming that in each case experiment, the order and pattern of construct destruction remained the same, regardless of data handling method.

Average cleavage profiles for separase biosensor experiments were produced in Fiji by creating a clipping mask to the DNA using the far red signal emitted by SiR DNA treatment. The eGFP and mCherry intensity readings from the clipping mask were then plotted over time and aligned at PB1 extrusion. eGFP/mCherry ratios were calculated in Excel.

When only two constructs are compared, we have displayed both the average trace and individual traces. Where 3 or more constructs are compared, for clarity, only the average trace is shown with SEM error bars.

## Acknowledgements

We thank Professor Jonathan MG Higgins for scientific discussion throughout this study and for critical reading of the manuscript along with Dr. Alexandre Webster. We thank F. Davidson for technical assistance. This work was supported by a Wellcome Trust Career Re-entry Fellowship grant to SM [062376]. ORD is a Sir Henry Dale Fellow jointly funded by the Wellcome Trust and Royal Society [Grant Number 104158/Z/14/Z].

## Author contributions

CT carried out all experiments alongside SM, with critical contribution from BH in western blotting experiments and MDL in microscopy. The project was carried out under the supervision of SM and MDL. CT and SM designed the experiments with assistance from ORD. CT and SM prepared the manuscript with assistance from ORD. The authors declare no competing financial interests.

